# Rapid, seamless generation of recombinant poxviruses using host-range and visual selection

**DOI:** 10.1101/467514

**Authors:** Sameera Vipat, Greg Brennan, Sherry L. Haller, Stefan Rothenburg

## Abstract

Vaccinia virus (VACV) was instrumental in eradicating variola virus (VARV), the causative agent of smallpox, from nature. Since this first use as a vaccine, VACV has been developed as a vector for therapeutic vaccines and as an oncolytic virus. These applications take advantage of VACV’s easily manipulated genetics and broad host range as an outstanding platform to generate recombinant therapeutics. Several methods have been developed to generate recombinant VACV, including marker selection methods and transient dominant selection. Here, we present a refinement of a host-range selection method coupled with visual identification. Our method takes advantage of selective pressure generated by the host antiviral protein kinase R (PKR) coupled with a fluorescent fusion gene expressing mCherry-tagged E3L, one of two VACV PKR antagonists. This method permits rapid, seamless generation of rVACV in a variety of cell types.

## Introduction

Vaccinia virus (VACV) was the virus instrumental for the first successful eradication of a human pathogen, variola virus (VARV), from nature. Ever since the extermination of variola virus, poxviruses including VACV have continued to be useful therapeutic viruses for both human and animal medicine. For example, a VACV-based rabies virus vaccine has been very effective in preventing transmission of sylvatic rabies in Europe (1) and the United States (2). More recently, recombinant poxviruses expressing a variety of pro- and anti-tumor molecules, for example, single-chain antibodies or human erythropoeitin, have seen encouraging success in clinical trials as oncolytic agents (3–5). VACV is particularly attractive as a vector because it is readily amenable to genetic manipulation, possesses a broad host range, and it is stable under a variety of conditions, allowing easy transportation and vaccine viability in the field (6, 7). While multiple techniques have been developed to generate recombinant VACV for laboratory experiments and vaccine generation, current strategies to generate these viruses have notable limitations.

Broadly, there are two primary approaches to generate recombinant poxviruses. The first strategy employs homologous recombination to introduce a cassette including the transgene and a selectable marker gene such as antibiotic resistance. The cassette is flanked by two ∼500nt or larger homology arms directing the gene to a specific site in the viral genome, which is then stably integrated by double crossover events (8–10). This strategy is rapid and efficient; however, it results in extra genetic material in the form of the marker gene that may produce unexpected effects. Furthermore, there is a practical upper limit to the number of transgenes that can be introduced limited by the number of unique selectable markers available. Transient dominant selection (TDS) strategies have addressed this issue by facilitating the generation of “scarless” recombinant viruses (11). Using this strategy, a plasmid containing a mutant VACV gene and a selectable marker gene are integrated into the viral genome, but without flanking VACV DNA. This approach results in transient integration of the entire plasmid and duplication of the VACV gene as a result of integration by a single crossover event. This intermediate is stable as long as it’s maintained under selection pressure, permitting enrichment of this construct. When selection is removed, the VACV duplication enables a second crossover event that results in the removal of the plasmid and subsequent formation of either the wildtype (wt) or recombinant virus in approximately 50:50 ratio. While TDS generates recombinant viruses without requiring the stable introduction of foreign DNA, resulting viruses must be screened for the expected mutation by sequencing analysis, a potentially time consuming and costly step.

Here, we present an approach to generating recombinant poxviruses combining the best aspects of each of these approaches. Our strategy combines visual and host-range selection to rapidly generate recombinant viruses by double crossover events, and subsequently eliminate the selectable marker gene by homologous recombination. This approach permits the rapid generation of mutants mediated by homologous recombination, with the “scarless” nature of TDS approaches while not requiring a subsequent screening step to distinguish wt and mutant viruses.

Our method also uses host-range selection in place of antibiotic selection, eliminating the risk of chemically induced phenotypic changes in the cell line. For this approach, we have chosen to use the host antiviral protein kinase R (PKR) as the selective agent to generate recombinant VACV. PKR is expressed as an inactive monomer in most cell types (12). Upon binding double-stranded RNA (dsRNA) at the N-terminal dsRNA-binding domains, PKR dimerizes and is autophosphorylated (13). This active form of PKR phosphorylates the α-subunit of the eukaryotic initiation factor 2 (eIF2α), ultimately inhibiting delivery of methionyl-tRNA to the ribosome, thereby preventing intracellular translation and broadly inhibiting the replication of most known virus families (14, 15).

In response to the broad and potent antiviral activity of PKR, many viruses have evolved at least one strategy to prevent PKR activation. Most poxviruses express two PKR antagonists, designated E3L and K3L in VACV, that antagonize PKR through two distinct mechanisms(16). E3 prevents PKR homodimerization by binding double-stranded RNA(17, 18), while K3 acts as a pseudosubstrate inhibitor by binding directly to activated PKR and thereby inhibiting interaction with its substrate eIF2α(19). Importantly, these two PKR antagonists do not necessarily inhibit PKR from all species. For example, the K3 homolog from sheeppox virus strongly inhibited PKR from sheep, whereas the sheeppox E3 homolog did not show PKR inhibition. Therefore, it is necessary to delete both E3L and K3L to ensure that VACV cannot replicate in cells from any species. In this study, we present a method to use PKR-mediated selective pressure combined with fluorescence selection to generate a VACV recombinant deleted for E3L and K3L (VC-R4) that cannot replicate in PKR competent cells derived from diverse species. This recombinant virus provides an excellent background for rapid generation of recombinant viruses expressing genes under control of the native E3L promoter.

## Materials

### Cell lines

European rabbit kidney cell line RK13 and RK13+E3L+K3L (RK13++, (20)) cells were maintained in Dulbecco’s Modified Essential Medium (DMEM, Life Technologies) supplemented with 5% FBS (Fisher) and 25 µg/ml gentamycin (Quality Biologicals). RK13++ cells were also maintained in 500 µg/ml geneticin and 300 µg/ml zeocin (Life Technologies).

### Viruses

VP872 (kindly provided by Dr. Bertram Jacobs) is derived from vaccinia virus (VACV) Copenhagen strain (VC2) and lacks the PKR-inhibitory gene K3L as described previously (19). All viruses were titered in RK13++ cells by serial dilution.

### Primers

SR1_5pE3L_F – 5’-gtaggaatactacgccgatg

SR2_5pE3L_R+msc – 5’-gagctcctcgagaagcttcaattgtcgacatttttagagagaactaacacaaccagc

SR3_p11_eGFP_F – 5’-gaagcttctcgaggagctcgtagaatttcattttgtttttttctatgctataaatggtg

SR4_EGFP_R – 5’-ctaaactactaactgttattgataactagaattacttgtacagctcgtccatg

SR5_E3L_3repeat_F – 5’-ttctagttatcaataacagttagtagtttag

SR6_E3L_3repeat_R – 5’-ttatattccaaaaaaaaaaaataaaatttcaatttttagacgaatatctgtgacagat

SR7_mCherry_F – 5’-ctaaaaattgaaattttattttttttttttggaatataaatggtgagcaagggcgagg

SR8_E3L_R – 5’-aactagaaggtaccttatcagaatctaatgatgacg

### Plasmids

– p688-mCherry-E3L

pD2EGFP-N1 (Clontech)

pS96, containing eGFP surrounded by 5’ and 3’ sequences homologous to the region flanking

VACV Copenhagen E3L (21)

## Methods

### Recombination plasmid construction

We constructed a recombination cassette containing 518 nucleotides homologous to the VACV genomic region 5’ of E3L (5’ homology arm), eGFP, 150 nucleotides homologous to the VACV genomic region 3’ of E3L (short 3’ homology arm), a mCherry-E3L fusion gene, and 567 nucleotides homologous to the VACV genomic region 3’ of E3L including the short 3’ homology arm (long 3’ homology arm, Fig. 1). Primers SR1 and SR2 amplified the 5’ homology arm, and SR5 and SR6 were used to amplify the short 3’ homology arm from pS96 (21), SR3 and SR4 were used to amplify eGFP from pD2EGFP-N1 (Clontech), and SR7 and SR8 were used to amplify the mCherry-E3L fusion from p688. Fragments were amplified in a 50 uL reaction volume composed of 10 μL Phusion HF buffer (5X), 1 μL dNTP (10 mM), 2.5 μL of each primer (10 μM), 0.5 μL of Phusion DNA polymerase, template DNA (10 ng), and nuclease free water to 50 uL. Amplicons were gel purified, and the concentration of each gel-purified amplicon was estimated by gel electrophoresis. Equimolar amounts of each of the four amplicons were combined by splicing by overhang extension PCR (SOE-PCR) in a 50 uL reaction volume composed of 10 μL Phusion HF buffer (5X), 1 μL dNTP (10 mM), 2.5 μL each of primers SR1 and SR8 (10 μM), 0.5 μL of Phusion DNA polymerase, template DNA (2 μL reaction 1, 1 μL reaction 2, 6 μL reaction 3, and 2 μL reaction 4), and nuclease free water to 50 uL. The SOE-PCR cycling protocol was initial denaturation at 98°C x 1 minute, 30 cycles of 98°C x 10 seconds, 52C x 30 seconds, 72°C x 90 seconds, a final extension of 72°C x 7 minutes. This amplicon was gel purified, and 5 μL of this product was incubated at 68°C for 7 minutes with 0.5 μL 10 mM dATP, 3 μL 5X OneTaq buffer, 0.5 L OneTaq polymerase, and 3 uL dH2O to add single(A) 3’ overhangs, then cloned into pCR2.1 using the TOPO TA Cloning Kit following the manufacturer’s protocol (ThermoFisher). TOP10 chemically competent *E. coli* were transformed with the resulting product and subjected to blue/white screening. The insert was verified by diagnostic endonuclease digestion with *EcoRI*, and confirmed by Sanger sequencing. Finally, the long 3’ homology arm was derived from the previously described plasmid S96 (21) which encodes eGFP flanked by 5’ and 3’ homology arms derived from VACV. Both pS96 and the p837-GOI-mCherry-E3L were digested with ClaI, present in the 5’ homology arm of both pS96 and p837-GOI-mCherry-E3L, and KpnI, immediately 5’ of the long 3’ homology arm in pS96 and present in primer SR8 3’ of the mCherry-E3L fusion. These restriction fragments were ligated in a 2:1 molar ratio using T4 ligase (NEB) at 4°C for 16 hours, and this product was transfected into chemically competent *E. coli*. We screened individual colonies for the appropriate insert by KpnI/ClaI digestion and by Sanger sequencing.

**Figure 1.**
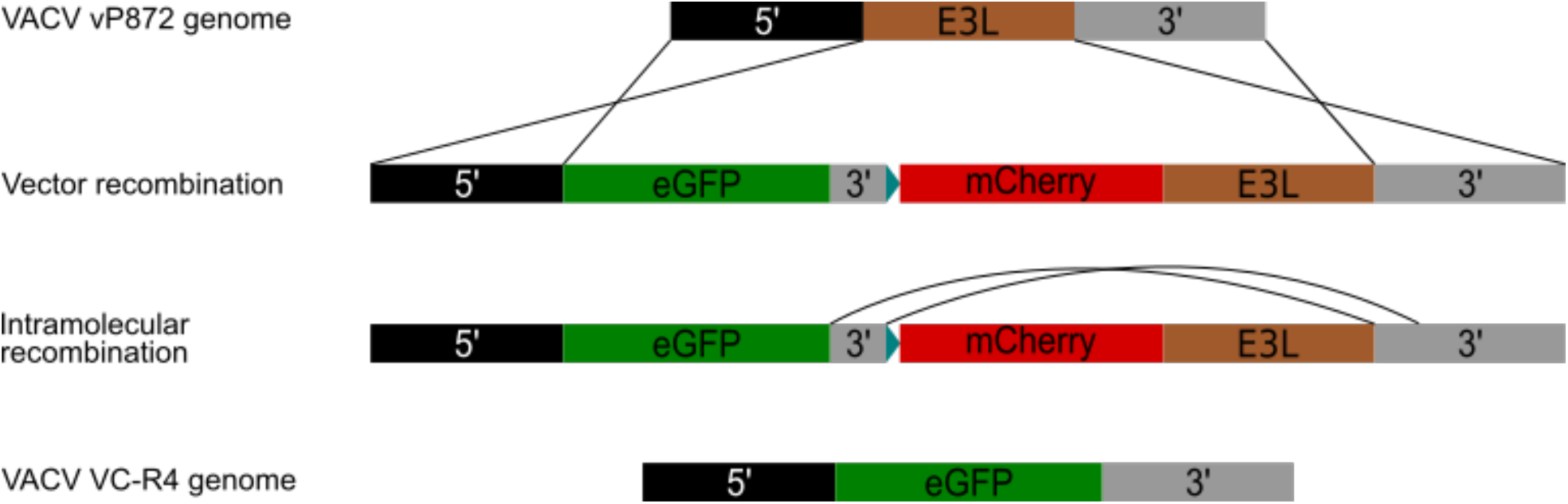
Diagram of p837-GOI-mCherry-E3L as well as the host-range and visual recombination strategy. **(A)** 5’ homology arm (black) and 3’ homology arm (grey) flank the E3L locus (brown) in VACV. **(B)** In p837-GOI-mCherry-E3L, these homology arms flank a cassette containing eGFP (green) separated from an mCherry-E3L (red) fusion gene under control of the synthetic early/late poxvirus promoter (26) blue) by a short 3’ homology arm (grey). These external homology arms drive homologous recombination between VACV and the p837-GOI-mCherry-E3L. **(C)** When PKR selective pressure is removed, intramolecular recombination occurs between the short and long 3’ homology arms. **(D)** Resulting in a virus (VC-R4) containing the gene of interest in the E3L locus, with a short, untranslated molecular tag resulting from the multiple cloning site in p837-GOI-mCherry-E3L.

This recombination cassette contains a 5’ SalI recognition sequence that cuts immediately 3’ of the native E3L start codon. Therefore, this site allows the gene of interest to be expressed under control of the endogenous E3L promoter. It should be noted that because of this strategy, subsequent genes should be subcloned into the eGFP locus without an initial start codon. Finally, we modified the cassette immediately 3’ of the eGFP locus to introduce three additional restriction sites for NdeI, NheI, and BamHI between eGFP and the long 3’ homology arm. These sites facilitate the subcloning of different genes of interest into this recombination vector (p837), and also act as a unique molecular identifier to label recombinant viruses.

### Generation of VC-R4

To ensure that our PKR-mediated host range selection was efficient, we first used this recombination plasmid to generate a recombinant VACV lacking both E3L and K3L. E3L and K3L have been shown to have overlapping but non-redundant PKR inhibitory activity in cells derived from different species {Park, 2019; (22, 23). Therefore, to insure a broad range of suitable cell types for this selective strategy, it was necessary to knock out both E3L and K3L to provide for the broadest range of cell types can be used for subsequent virus selection. To this end, we selected vP872, a virus already lacking K3L, as the background for our E3L knockout.

We infected confluent 6-well plates of wildtype European rabbit kidney cell line RK13 with VP872 (MOI = 1.0). One hour post-infection, we replaced the infecting medium with fresh DMEM, and transfected these cells with 500 ng of p837-GOI-mCherry-E3L using Lipofectamine 2000 following the manufacturer’s protocol (ThermoFisher). PKR-mediated selective pressure maintained expression of the mCherry-E3L fusion protein in these cells to permit replication of the recombinant virus. We identified recombinant viruses by fluorescence microscopy, as plaques from these viruses expressed both red fluorescence due to integration the mCherry-E3L fusion gene and green fluorescence from the eGFP gene in p837-GOI-mCherry-E3L (Fig. 2). We plaque purified viruses exhibiting both red and green fluorescence were on wt RK13 cells three times. To permit collapse of the mCherry-E3L selection marker for “scarless” generation of VACV*Δ*E3L*Δ*K3L+eGFP (VC-R4), we infected RK13 RK13+E3L+K3L cells (20). These cells provide the VACV PKR antagonists in *trans* and alleviate the PKR-mediated selective pressure to maintain the mCherry-E3L fusion gene. Because this fusion gene is flanked by the short 3’ homology arm and the long 3’ homology arm in close proximity, intramolecular homologous recombination is favored and the fusion gene is lost relatively rapidly. Viruses that have undergone this recombination step are identified by eGFP-positive, mCherry-negative plaques by fluorescent microscopy. These single-positive plaques were subjected to three rounds of plaque purification in RK13+E3L+K3L cells. We purified DNA from the plaque purified virus as previously described (24) and confirmed that this virus lacked both E3L and K3L by Sanger sequencing. This virus is named VC-R4.

**Figure 2.**
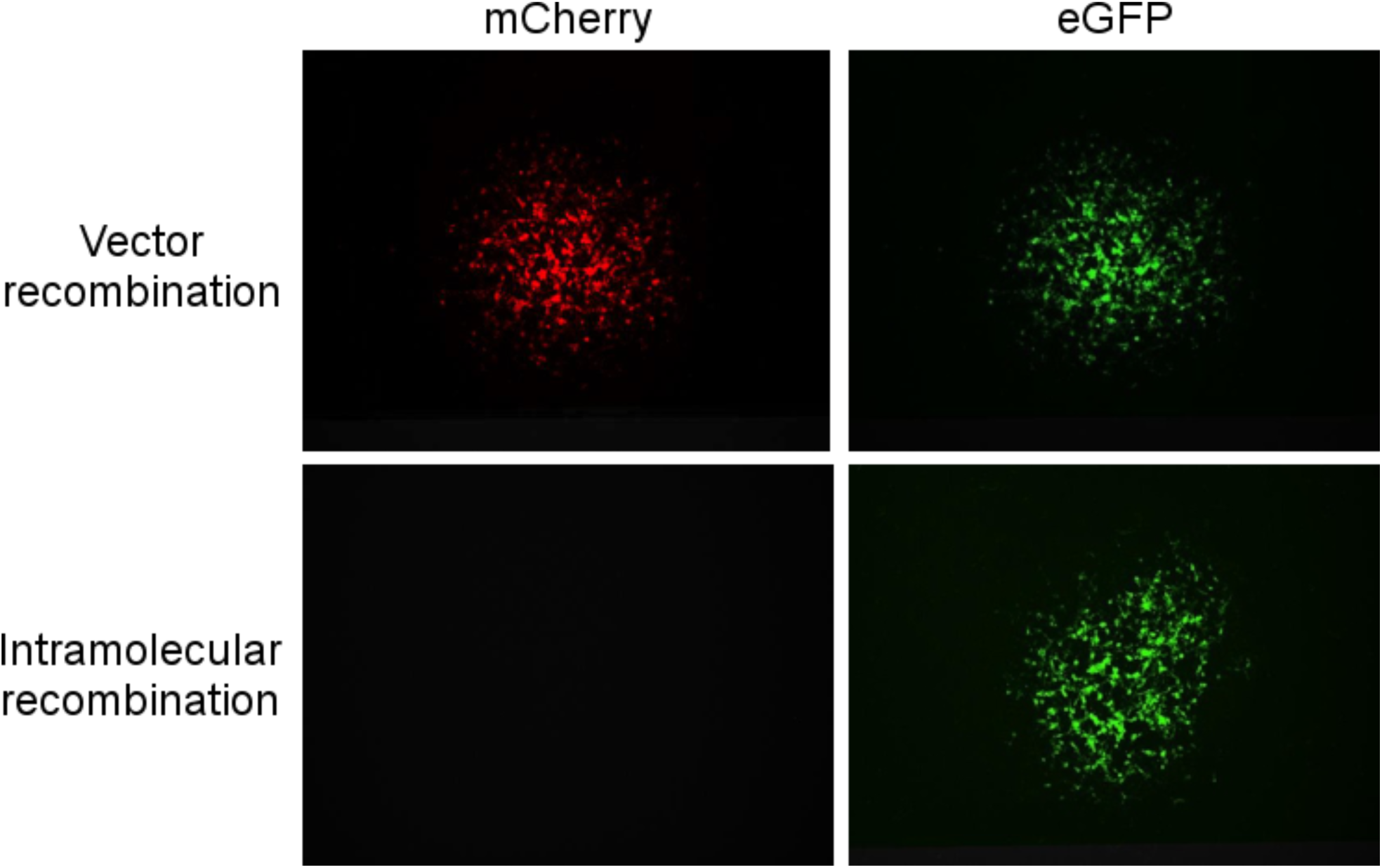
Fluorescent micrographs of (top) a recombinant virus plaque after recombination with p837-GOI-mCherry-E3L expressing both EGFP (left) and mCherry (right) in RK13 cells. (Bottom) Micrograph of a recombinant virus plaque after PKR-mediated selective pressure has been removed in RK13++ cells, expressing mCherry (right) but not EGFP (left).

### VC-R4 does not replicate in PKR-competent cells

To determine how the loss of both VACV PKR antagonists, E3L and K3L, affected VC-R4 replication, we titered VC-R4 and vP872 on multiple cell types by serial dilution (Fig. 3). As expected, vP872 replicated to substantially similar titers in PKR competent RK13 and HeLa cell lines, while we did not detect any plaques in VC-R4 infected RK13 or HeLa cells. In both of these cells, eIF2α is phosphorylated after VC-R4 infection, consistent with PKR activation inhibiting VC-R4 replication {Park, 2019}. However, in RK13+E3L+K3L cells, the expression of these VACV PKR antagonists in *trans* rescued VC-R4 replication, although VC-R4 replicated 4-fold less efficiently in the complimenting cell line relative to vP872. We have observed this phenotype in other replication deficient viruses infecting RK13+EL+K3L, and hypothesize that this incomplete rescue is most likely due to E3 and K3 being supplemented in trans and therefore not necessarily delivered to the VACV replication factories with the appropriate timing or concentration to fully inhibit PKR activation. Taken together, these data demonstrate that VC-R4 is potently restricted by PKR activation in cells derived from multiple species.

**Figure 3.**
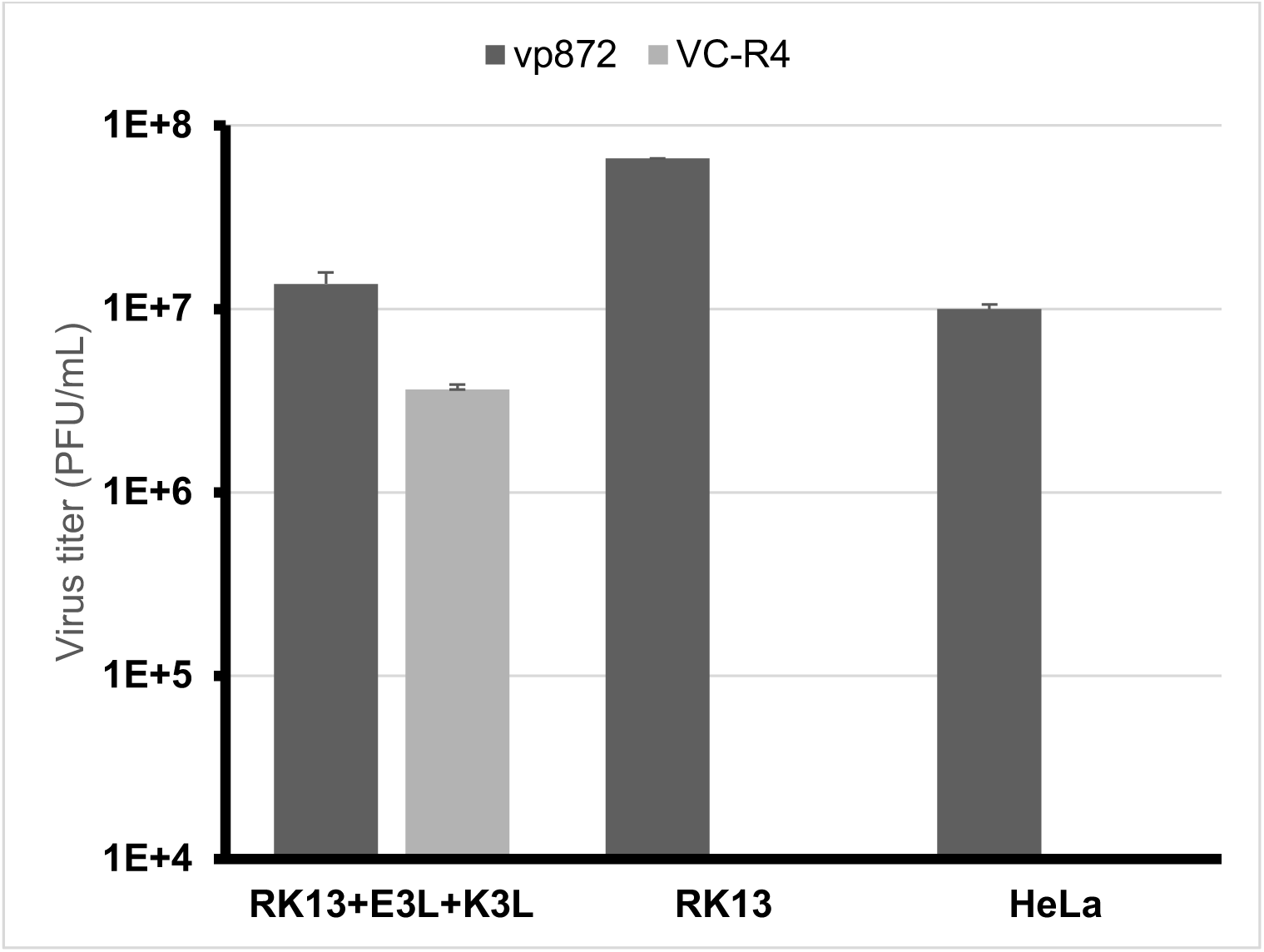
VC-R4 cannot replicate in PKR competent cells. The indicated cell lines were infected with vP872 (dark grey) and VC-R4 (light grey) (MOI = 0.1) 48 hours post-infection the infected cells were harvested and titered by serial dilution on RK13++ cells. Titers are reported in PFU/mL.

## Discussion

Here we present a variation of a transient marker selection strategy (25) to generate recombinant vaccinia viruses without introducing foreign DNA in the final recombinant virus. Our strategy uses selective pressure mediated by the host antiviral protein PKR rather than other forms of selective pressure such as antibiotics. The use of host antiviral genes eliminates the possibility of chemically induced phenotypic changes in the cells, or increased risk of mutation due to selection drugs. Furthermore, unlike with drug selection, there is no lag phase for our approach, because PKR is expressed constitutively in nearly all cells. Secondary visual selection based on mCherry expression also improves the specificity of this method by ensuring that only plaques expressing the transgene are picked during the first phase, and is equally efficient as a negative selective marker while selecting mature recombinant viruses that have lost the mCherry-E3L gene.

Here, we demonstrate the use of this method to generate a VACV recombinant deleted for both PKR antagonists E3L and K3L and expressing eGFP under control of the E3L promoter. Going forward, this virus will serve as an efficient background for future recombinant viruses, as it is incapable of replicating in PKR competent cells. Therefore, there will be strong PKR-mediated selective pressure to drive the mCherry-E3L recombination cassette into progeny virions while at the same time essentially preventing replication of non-recombinant virus. Furthermore, the loss of EGFP by uptake of the recombination cassette is a useful secondary selection marker to ensure picked plaques are not co-infected with a non-recombinant virus. As demonstrated by the insertion of EGFP, with this approach, any gene can be rapidly inserted into the E3L locus under control of the native promoter, provided that PKR null cells or a complimenting cell lines are used for downstream experiments if the transgene is not a PKR antagonist. This strategy, combined with the VC-R4 virus that we report here, adds a new and potent method to rapidly and reliably generate recombinant vaccinia viruses using host-mediated selective pressure and visual identification of recombinants early in the process.

